# A decision underlies phototaxis in an insect

**DOI:** 10.1101/023846

**Authors:** E. Axel Gorostiza, Julien Colomb, Björn Brembs

## Abstract

Like a moth into the flame - Phototaxis is an iconic example for innate preferences. Such preferences likely reflect evolutionary adaptations to predictable situations and have traditionally been conceptualized as hard-wired stimulus-response links. Perhaps therefore, the century-old discovery of flexibility in *Drosophila* phototaxis has received little attention. Here we report that across several different behavioral tests, light/dark preference tested in walking is dependent on various aspects of flight. If we temporarily compromise flying ability, walking photopreference reverses concomitantly. Neuronal activity in circuits expressing dopamine and octopamine, respectively, plays a differential role in photopreference, suggesting a potential involvement of these biogenic amines in this case of behavioral flexibility. We conclude that flies monitor their ability to fly, and that flying ability exerts a fundamental effect on action selection in *Drosophila*. This work suggests that even behaviors which appear simple and hard-wired comprise a value-driven decision-making stage, negotiating the external situation with the animal’s internal state, before an action is selected.

## Introduction

In their struggle for survival, animals need not just the capability to trigger behaviors at the appropriate time, but these behaviors need to be flexible in response to or anticipation of changes in environmental and internal conditions. What may be an appropriate response to a given stimulus when the animal is hungry may be maladaptive when the animal is seeking a mating partner, and *vice versa*. The relative values of extrinsic and intrinsic factors must be analyzed and weighed in order to shape the behavior to be adaptive in a particular situation. Across animal phyla, biogenic amines have been found to be part of a complex network involved in such value-driven processes. In invertebrates, Dopamine (DA) and Octopamine (OA) are two important modulators of behavior. OA, the invertebrate counterpart of the adrenergic vertebrate system, has been implicated in state-dependent changes in visual-processing [1,2], experience-dependent modulation of aggression [3], social decision-making [4], and reward [5]. DA is also known for its countless roles in physiological and behavioral processes across animal phyla such as reward [5–7], motivation [8–10] and value-based or goal-directed decision-making [8,11–15]. Complementing such flexible behaviors are simple, innate responses such as escape responses, taxis/kinesis behaviors, or fixed action patterns. They are commonly thought to be less flexible and more automatic, but with the advantage of either being especially efficient, fast, or with only a low cognitive demand. However, recent research has shown that many of these behaviors are either more complex than initially imagined [16–19] or liable to exploitation [20]. Moreover, several studies have shown that the state of the animal modulates how sensory structures process identical stimuli [21–26] and many of these modulations are caused by aminergic actions [1,2,21,27–29]. Due to observations like these, the general concept of behaviors as responses to external stimuli (‘sensorimotor hypothesis’) has come under ever more critical scrutiny in the last decade. Studying what can arguably be perceived as the most iconic of stereotypic insect responses, the approach of a bright light (phototaxis), we provide further evidence that the simple input-output relationships long assumed to underlie most if not all behaviors, may only exist at the observational level, dissipating at the neuronal level.

*Drosophila melanogaster* phototactic behavior has been studied for at least one hundred years. As most flying insects, flies move towards a light source after being startled, showing positive phototaxis. This innate preference for light appears to be species- and strain-specific and has been described as part of a fly’s personality [30]. Recently, it has been shown that mated female flies transiently avoid UV light during egg-laying [31]. Interestingly, experiments described by McEwen in 1918 and Benzer in 1967 demonstrated that wing defects affect phototaxis also in walking flies. These early works showed that flies with clipped wings did not display the phototactic response to light, whereas cutting the wings from mutants with deformed wings did not decrease their already low response to light any further [32,33]. The fact that manipulating an unrelated organ, such as wings, affects phototaxis contradicts the assumed hard-wired organization of this behavior, suggesting that it may not be a simple matter of stimulus and rigid, innate response, but that it contains at least a certain element of flexibility. In this work, we systematically address the factors involved in this behavioral flexibility and begin to explore the neurobiological mechanisms behind it.

## Methods

### Strains and fly rearing

Flies were reared and maintained at 25°C in vials containing standard cornmeal agar medium [34] under 12h light/dark cycles with 60% humidity, except for experiments involving *UAS-trpA1* or *UAS-shibire*^*TS*^, in which parental crosses and their offspring were maintained at 18°C under 12h light/dark cycles with 60% humidity.

Stocks obtained from the Bloomington Drosophila Stock Center (RRID:SCR_006457; NIH P40OD018537) were used in this study: *UAS-TrpA1* (26263), th-GAL4 (8848), tdc2-GAL4 (9313), and *PKC*^*δ*^ (18258). The *PKC*^*δ*^ mutant flies were intended for a different project when we discovered that the flies do not even attempt to fly. To our knowledge, the molecular mechanism behind the flightlessness is unknown.

The sources of other stocks are detailed here:

*w*^1118^, *w*^1118^; *hs*-Gal4 (heat shock inducible GAL4), and *UAS*-*PKCi* (inhibitory pseudosubstrate of protein kinase C) were provided by Henrike Scholz (University of Cologne, Germany).

*WTB* is a Wild-type Berlin strain from our stock in Regensburg.

*CS*^*RE*^ is a *Canton S* strain bred in our lab in Regensburg.

*CS*^*TZ*^ and *FoxP*^*3955*^ were provided by Troy Zars (University of Missouri, USA).

*rsh*^*1*^ was provided by B. van Swinderen (The University of Queensland, Australia).

*rut*^*2080*^, *mb247*-GAL4 and *UAS-CNT-E* were provided by Martin Heisenberg (Rudolf Virchow Center, Germany).

*act88F-Gal4* was provided by Juan A. Navarro (University of Regensburg, Germany).

*A9*-GAL4 and *UAS-baboon*^*QD*^ were provided by Florian Bayersdorfer (University of Regensburg, Germany).

### Mechanical manipulations

Unless described otherwise, 24h before the experiment 2-5 d old flies were briefly anesthetized under CO_2_. In the standard wing-clipping procedure, the distal two thirds from both wings were clipped from half of the individuals (Fig. 1A). At least 30 flies with clipped wings and 30 flies with intact wings were placed in the same vial until the experiment was performed, in which they were tested together. For other manipulations, one of the different treatments (see Fig. 1) was applied to half of the flies of a given group. At least sixty flies (half of them with injury) were placed in vials for a 24h recovery period and tested together. Flies with abdominal injury were not mixed with intact flies to avoid mistakes during the evaluation of the experiment due to the inconspicuous nature of the injury.

**Figure 1.**
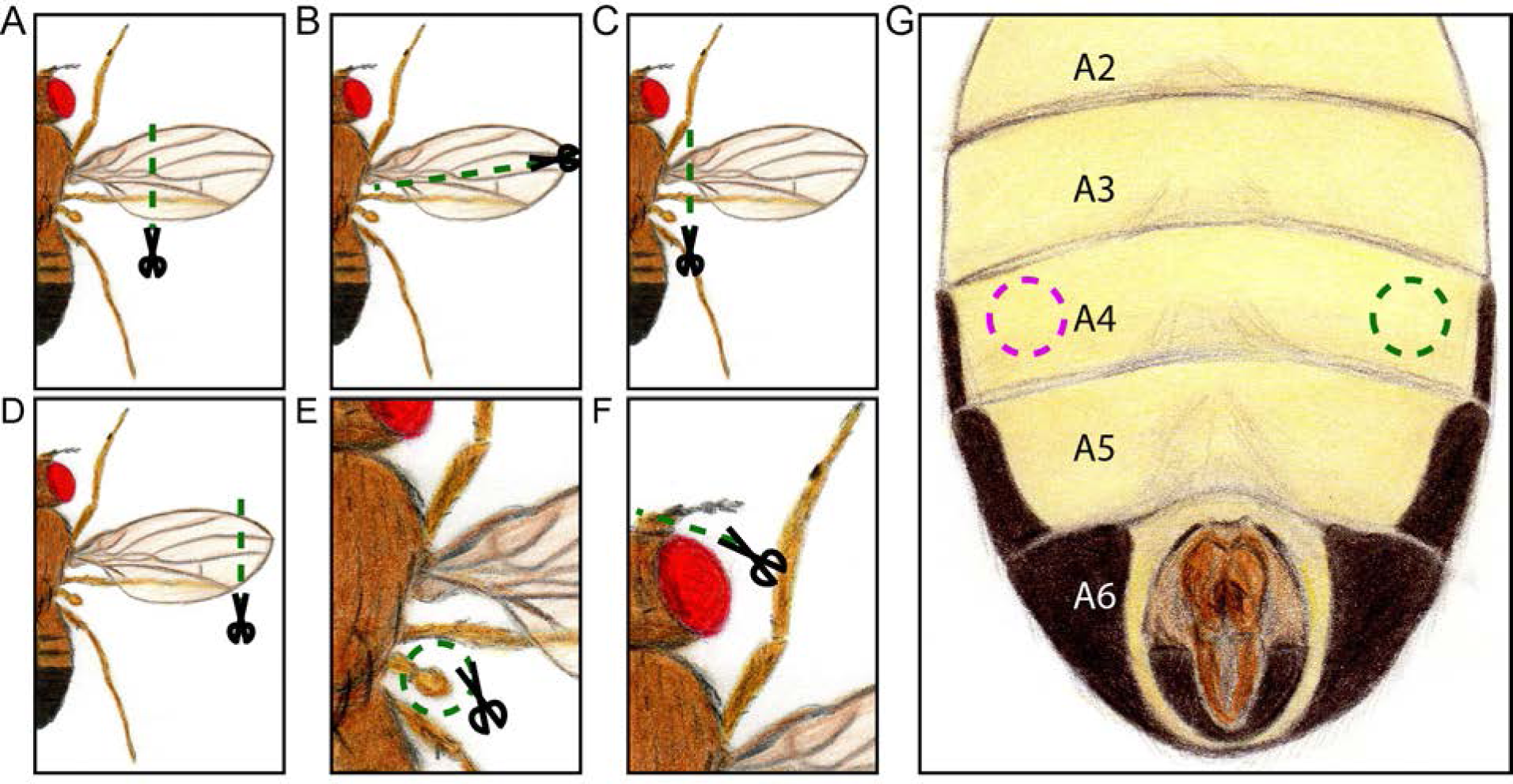
Schematic representation of the different injuries made to the flies. ***A***, This was the standard procedure, where the distal two thirds from both wings were removed. ***B***, Longitudinal cut. Half of the wing was removed. It was applied to both wings in experiments of Fig. 4A,B. ***C***, Whole wing cut. It was used in Fig. 4C,D to remove only one wing (the side was randomly selected), and in Fig. 4E,F to remove both wings. ***D***, End of the wing cut. Around 20% of each wing was removed. It was used in Fig. 4E,F. ***E***, Haltere removal. Both halteres were removed and the effect on photopreference is presented in Fig. 4G,H. ***F***, Antennal damage. The third segment of both antennae was cut. This treatment was used for experiments in Fig. 4I,J. ***G***, Abdominal injury. Flies were stabbed on one side of the ventral fourth abdominal segment (the side was randomly selected). The results of the effect of this injury in phototaxis are depicted in Fig. 4K,L.

Haltere removal was performed by pulling each haltere with forceps, while the antennal damage was produced by clipping the third segment of the antenna (funiculus). The abdominal injury was performed with a sharpened needle, and was always made ventrally in one side of the fourth abdominal segment.

### Wing gluing

Flies were cold anesthetized using a custom made cold air station and their wings were glued together in their natural relaxed posture using a 3M sucrose solution. To unglue the wings flies were cold anesthetized and their abdomen gently submerged in water to dissolve the sucrose. After each process flies were left to recover overnight. Flies were discarded from the analysis if their wings were damaged because of the treatments or unglued by chance.

### Countercurrent Apparatus

Phototactic preference was evaluated using Benzer’s classic countercurrent apparatus [32] (http://dx.doi.org/10.17504/protocols.io.c8gztv). The apparatus was completely transparent and consisted of two acrylic parts, a lower one with 6 parallel tubes (an initial tube + 5), and a movable upper part with 5 parallel test tubes. Each plastic tube had a length of 6.8 cm, an inner diameter of 1.5 cm, and an outer diameter of 1.7 cm. The test group was placed in the initial tube and was left in darkness to acclimate for 10 min, with the apparatus placed horizontally. Thereafter, flies were startled by tapping the apparatus, making all of them end up at the bottom of the tube. The apparatus was placed horizontally and the upper part shifted, making the initial tube face the first test tube for 15 seconds, allowing the flies to move towards the light if the test tube was facing it (positive phototaxis test), or away from it if the initial tube was facing the light (negative phototaxis test). Then, the upper part was shifted again and flies that moved to the test tube were transferred to the next tube of the lower part by tapping the apparatus, and the same test was repeated 4 more times. The light source was always placed at 30 cm from the apparatus and consisted of a fluorescent warm white tube (OSRAM 18W/827), which delivers 1340 lux at that distance.

The Performance Index was calculated using the formula:

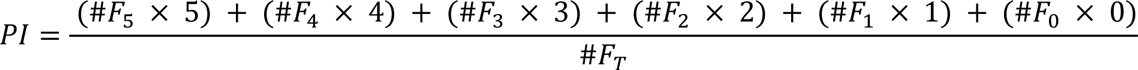

where *#F*_*n*_ was the number of flies in the tube n (being 0 the initial tube and 5 the last test tube), and *#F*_*T*_ was the total number of flies. If the test tubes were on the bright side a higher index meant a more positive phototaxis. In each experiment a PI was calculated for the wingless flies and other for the intact flies. The tubes were cleaned thoroughly after each test.

In order to facilitate comparisons in figures 3A and 6A, the effect size was calculated using the Glass Δ estimator.

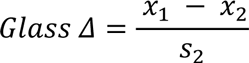

where *x*_1_ was the mean of treated group, *x*_2_ the mean of the control group, and *s*_2_ the standard deviation of the control group. When positive phototaxis was tested, a negative *Glass Δ* value reflected a reduction in positive phototaxis after wing-clipping; and when negative phototaxis was tested, a positive value represented an increase in negative phototaxis after wing-clipping.

### T-Maze

Light/Darkness choice was measured in a custom built, opaque PVC T-Maze with only one transparent (acrylic) choice tube (http://dx.doi.org/10.17504/protocols.io.c8azsd). Flies were placed in an initial dark tube (10 cm long, 1.5 cm inner diameter, and 2.5 cm outer diameter) and were left to dark adapt for 10 min. Then, they were transferred to the cylindrical elevator chamber (1.5 cm diameter, 1.5 cm height) by gently tapping the apparatus, where they remained for 30s. Next, the elevator was placed between the dark and the bright tube (both 20 cm long, 1.5 cm inner diameter, and 2.5 cm outer diameter), and flies were allowed to choose for 30s. As the source of light, the same fluorescent tube as for Benzer’s Countercurrent Apparatus was used, and placed 31.5 cm above the base of the T-Maze.

The Choice Index was calculated using the formula:

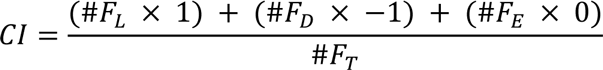

where *#F*_*L*_ meant the number of flies in the transparent tube, *#F*_*d*_ was the number of flies in the opaque tube, and *#F*_*E*_ was the number of flies that remained in the elevator. A *CI* of 1 meant all the flies chose the light, while an index of -1 meant a dark photopreference. The tubes were cleaned thoroughly after each round.

### Buridan

Locomotion towards dark objects was evaluated using Buridan’s paradigm as explained in Colomb *et al*. [35]. Briefly, 3-6d old flies were selected and half of them had their wings clipped under CO_2_ anesthesia (http://dx.doi.org/10.17504/protocols.io.c7vzn5). They were left to recover overnight within individual containers, with access to water and sugar (local store) before being transferred to the experimental setup. The setup consists of a round platform (117 mm in diameter) surrounded by a water-filled moat placed at the bottom of a uniformly illuminated white cylinder (313 mm in height) with 2 stripes of black cardboard (30mm wide, 313 mm high and 1 mm thick) placed 148.5 cm from the platform center one in front of the other. Flies were prevented from escaping by a transparent lid over the platform. The experiment duration was set to 900 seconds. Data were analyzed using BuriTrack and CeTrAn [35] (RRID:SCR_006331), both available at http://buridan.sourceforge.net.

### Genetic manipulation of wing utility and neuronal activity

For the experiments involving *TrpA1 and the act88f-GAL4* driver, experimental flies and their respective controls were raised at 18°C. 3-5d old flies were tested at room temperature (RT) and recovered for 5-6h at 18°C. Then, they were transferred to a 37°C climate room where they were placed in an acclimation vial for 15min. Next they were transferred to the first tube of the T-maze placed in the 37°C climate room, and the experiment proceeded as explained above. The choice step was reduced to 15s to compensate for the increased activity that flies showed in pilot experiments. After counting the flies, they were transferred to fresh vials and placed at 18°C for 24h. After this recovery phase, they were tested again at RT. We noticed that the CI obtained for wild types could differ between chambers at 37°C.

In the case of manipulation of dopaminergic and octopaminergic neural activity with *shi*^*TS*^ or *TrpA1* the same protocol was applied but instead of 37°C, 32°C were used and the choice step was 30s long.

### Statistical Analysis

Statistical analyses were performed with InfoStat, version 2013 (Grupo InfoStat, Facultad de Ciencias Agropecuarias, Universidad Nacional de Córdoba, Córdoba, Argentina) and R (http://www.r-project.org/). Number of replicates in each experiment was adjusted to provide a statistical power of at least 80% using pilot experiments. As dictated by the experimental design and data composition, a paired T-test, a Randomized Block Design ANOVA or an ANOVA were performed. Normality was tested using Shapiro-Wilks test, and the homogeneity of variance was assessed with Levene’s test. A value of p<0.05 was considered statistically significant. After ANOVA, a Tukey least-significant difference or an orthogonal contrasts test was performed. If an interaction between factors was significant in two-way ANOVAs, simple effects were performed, and p values were informed. In figures 1A, 3B-E and H, and 7C and D, homogeneity of variance was violated. In figures 1A, and 3B-E and H a Wilcoxon test was used, while in figures 7C and D Kruskal-Wallis test was employed for multiple comparisons. The alpha value was corrected using Bonferroni’s correction.

### Availability of data and materials

The datasets supporting the conclusions of this article are available in the FigShare repository, http://dx.doi.org/10.6084/m9.figshare.1502427.

## Results

### Wing-clipping effect is absent in flightless flies

Motivated by the findings of McEwen and Benzer, we decided to explore the nature of the phototactic change observed in wingless flies. After replicating Seymour Benzer’s original results on wild type flies and mutant flies with deformed wings (Fig. 2A), we wondered if the wing-clipping effect on phototaxis could be also observed in other genetic backgrounds. Therefore, flies with and without wings from two Canton-S strains inbred in different laboratories (*CS*^*TZ*^ *and CS*^*RE*^) and from the Wild Type Berlin (*WTB*) line were tested in Benzer’s Countercurrent Paradigm (BCP). All three lines showed a significant reduction in BCP performance index (PI) when the wings were cut (Fig. 2B). This reduction was apparent despite large variations between the three lines in the PI levels from intact flies, showing that the reduction in phototaxis due to wing-clipping can be observed across laboratory strains, with its magnitude dependent on genetic background and/or associated differences in baseline levels of phototactic performance.

**Figure 2.**
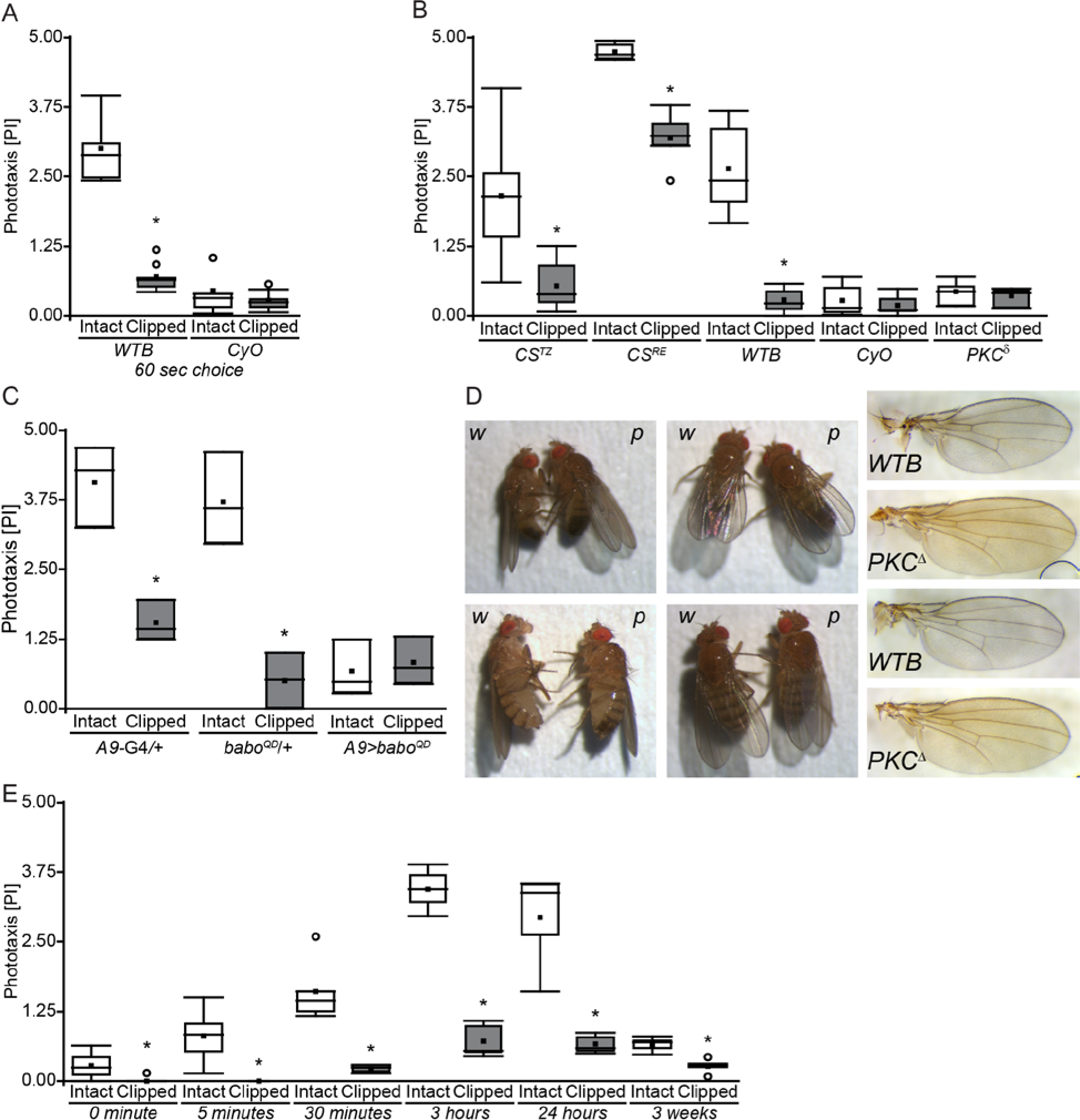
The wing-clipping effect is observable across genetic backgrounds and throughout adult lifespan, but is absent in flightless flies. ***A***, Replication of the original BCP experiments using 60s of time in which the animals were allowed to walk towards the light. Wilcoxon test; *WTB*: N=8, p<0.001; *CyO*: N=8, p=0.505 ***B***, BCP Performance Index (15s choice time) from three wild type strains and two flightless mutants with intact and clipped wings. Paired T-test; *CS*^*TZ*^: N=6, p=0.003; *CS*^*RE*^: N=5, p<0.001; WTB: N=12, p<0.001; CyO: N=14, p=0.066; *PKC*^*δ*^: N=4, p=0.413. ***C***, BCP Performance Index from flies with a genetic manipulation of wing development *(A9>babo*^*QD*^) and their genetic control groups (A9-G4/+, *babo*^*QD*^/+). Randomized Block Design ANOVA; N=3; Block p<0.001, Interaction Genotype vs Wings Integrity: p<0.001, simple effect Genotype: A9-G4/+: p<0.001, *babo*^*QD*^/+: p<0.001, *A9>babo*^*QD*^: p=0.401. ***D***, Lateral and dorsal view of wing posture of *WTB* (w) and *PKC*^*δ*^ (p) males (upper panels) and females (lower panels). Right panels: Examples of wing anatomy from *WTB* flies and *PKC*^*δ*^ mutant flies. *E*, BCP Performance Index of WTB flies after different recovery time lengths. Paired T-Test, 0 minutes: N=6, p=0.023; 5 minutes: N=6, p=0.008; 30 minutes: N=5, p=0.007; 3 hours: N=5, p<0.001; 24 hours: N=5, p=0.005; 3 weeks: N=5, p=0.004. * indicates significant differences. Box plot show quantiles 0.05, 0.25, 0.75 and 0.95, median, mean (black square), and outliers (circle).

Original experiments from McEwen, and then Benzer, showed that mutant flies with deformed wings displayed a lower positive phototaxis than wild types [32,33] and a diminished wing-clipping effect [33] (replicated in Fig. 2A). We wondered whether this simultaneous low phototaxis and absence of wing-clipping effect was due to a specific effect of these mutations or a general consequence of both manipulations altering the flies’ wing utility. In order to tackle this question, we tested three lines with flight impairments, the flightless *PKC*^*δ*^ mutant, the wings of which are indistinguishable from wild type wings (Fig. 2D), the *CyO* balancer line with curly wings, and a transgenic line in which the wings were deformed due to an overexpression of a constitutively active form of the *baboon* receptor in wing imaginal discs (*A9>babo*^*QD*^, [36]). Again replicating previous experiments, *CyO* flies showed a reduced PI that remained unchanged in wing-clipped animals (Fig. 2B). Similarly, *A9>babo*^*QD*^ showed less attraction to light and no significant wing-clipping effect (Fig. 2C), while all genetic controls behaved similar to wild type flies. Remarkably, *PKC*^*δ*^ mutants exhibited the same behavioral characteristics as *CyO* flies (Fig. 2B). Hence, we conclude that the reduction in phototaxis is not dependent on the origin of wing damage or the damage itself, but probably on wing utility.

### The behavioral change is immediate

If flies were able to assess wing utility, wing-clipping might have an almost instantaneous effect on the behavior. Thus, to find out when the behavioral change takes place, we assessed wing-clipped *WTB* flies at different time points after the injury was made. Flies from different groups were tested either 3 weeks, 24h, 3h, 30min, 5min or immediately after the surgery. To diminish the effects of anesthesia on phototactic behavior [37], we only used CO_2_ anesthesia for recovery times longer than 30min, and cold anesthesia for 0 and 5min recoveries. We found that the reduction in phototaxis could be observed in all tested groups (Fig. 2E). Moreover, the difference between intact and clipped flies increased with longer recovery phases, probably due to the vanishing of the anesthesia effect, only to decrease again in aged flies, perhaps due to a combination of a deteriorated locomotor activity and a decreased response to light in old flies [38,39]. Even if flies were placed in BCP right after surgery and let to recover from anesthesia only during the acclimation phase (0min group), it was possible to see a significant decrease in phototaxis. These results are consistent with the hypothesis that flies continually (or at relatively short intervals) monitor their ability to fly.

### Wingless and untreated flies do not differ in their locomotor activity

A potential explanation for the reduction in phototaxis is a possible reduction in locomotor activity in treated flies. We tested this hypothesis by placing the light source not only in front of the horizontal tubes of the BCP, but also above them, with the light shining perpendicular to the trajectory of the flies. In addition, we tested for negative phototaxis by placing the light source on the same side of the starting tube, such that we were able to count the flies with negative phototaxis. This tripartite experimental design allowed us to directly compare all three situations: light source on the opposite side of the starting tube (positive phototaxis), light source on top of the BCP (no taxis; locomotor activity control), and light source on the same side as the starting tube (negative phototaxis). In order to facilitate direct comparison of the behavioral consequences of wing-clipping in the three situations, we assessed the proportion of behavioral change with the *Glass Δ Effect Size* (ES). A negative ES in positive phototaxis indicates a reduction in positive phototaxis after wing-clipping. A negative ES in the no-taxis situation indicates a decrease in locomotor activity after wing-clipping, a positive ES an increase. A positive ES in the negative phototaxis situation indicates an increase in negative phototaxis after wing-clipping. We could not find any evidence for a reduced locomotor activity in these experiments. If anything, there was a small tendency of wing-clipped flies, instead of reducing their locomotor activity to actively avoid the light source (Fig. 3A).

**Figure 3.**
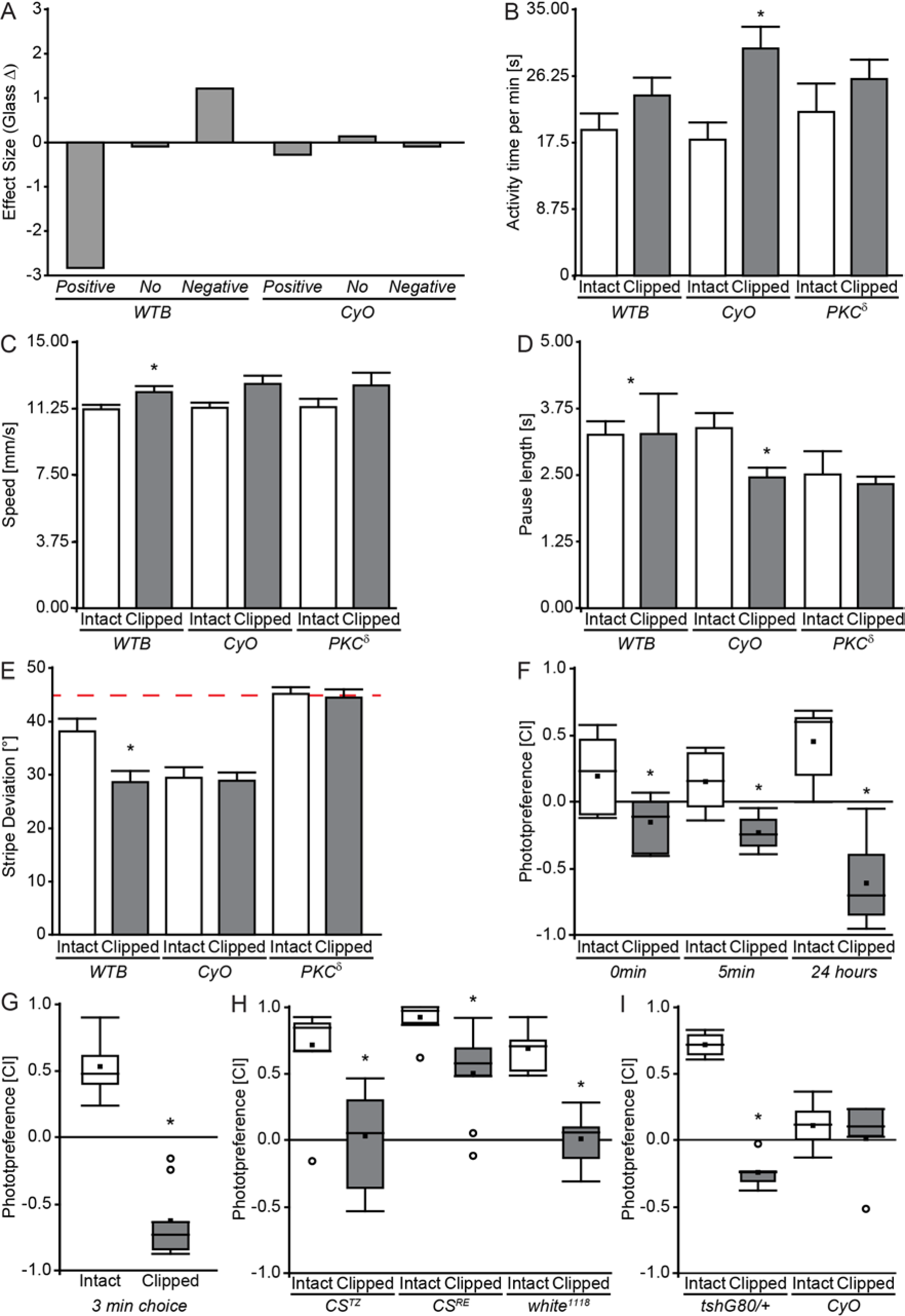
Flies without wings are not less active and prefer darker stimuli. **A**, Effect Size of wing clipping on BCP with the light source on the opposite side of the starting tube (positive phototaxis-*Positive-)*, light source on top of the BCP (no taxis *-No-;* locomotor activity control), and light source on the same side as the starting tube (negative phototaxis *-Negative-). B-E*, Buridan’s paradigm. *WTB*: Intact, N=20; Clipped, N=21. CyO, N=17. *PKC*^*δ*^, N=13. Wilcoxon test. *B*, Activity time. WTB: p=0.151. CyO, p=0.002. *PKC*^*δ*^, p=0.526. **C**, Speed. WTB: p=0.033. CyO, p=0.056. *PKC*^*δ*^, p=0.159. **D**, Pause Length. WTB: p=0.022. CyO, p=0.002. *PKC*^*δ*^, p=0.426. **E**, Stripe deviation. WTB: p=0.004. CyO, p=0.959. *PKC*^*δ*^, p=0.98. Dotted line indicates 45°, the mean value for computer-generated data. *F*, T-Maze Choice Index after different recovery time lengths. Paired T-Test; WTB: 0 minutes: N=7, p=0.003; 5 minutes: N=6, p=0.026; 24 hours: N=6, p<0.001. *G*, T-Maze Choice Index with 3 min choice step. Paired T-test; WTB: N=8, p<0.001. *H*, Choice Index of *CS*^*TZ*^, *CS*^*RE*^ and *w*^*1118*^ flies with intact and clipped wings. Wilcoxon test. *CS*^*TZ*^, N=8, p=0.003. *CS*^*RE*^, N=11, p<0.001. *w*^*1118*^, N=8, p<0.001. *I*, Choice Index of *CyO* flies and their wild-type siblings. Two way ANOVA, N=5, Interaction Wings Integrity (intact or clipped) vs Genotype p<0.001, simple effects: clipped vs intact: *tshG80/+* p<0.001, *CyO* p=0.487. See figure 2 for detailed graph information.

We tested the generality of these results in two additional experiments, Buridan’s paradigm and a T-maze. Buridan’s Paradigm, where the flies walk on a water-surrounded circular platform with two opposing vertical black stripes on the walls of a round panorama illuminated in bright white light from behind, has been used as a standard test for walking speed and locomotor activity for several decades [35,40]. We compared total activity time, walking speed, and pause duration in intact and wingless flies from three lines (*WTB*, *CyO*, *PKC*^δ^) in a modified version of Buridan’s Paradigm, where a roof prevents the flies from escaping. The results show only occasional small differences with the overall tendency of wingless flies exhibiting, if anything, slightly higher general activity than intact flies (Fig. 3B, C, D).

### Black stripe fixation in Buridan’s Paradigm is influenced by wing utility

Interestingly, the wing-clipped wild type flies also showed a stronger fixation of the black stripes in Buridan’s Paradigm, compared to the intact flies, while the flightless flies did not show such a difference (Fig.3E). This result is consistent with the tendency of the wild type flies to show some negative phototaxis after wing clipping (Fig. 3A). One possible explanation for these two congruent observations in such disparate experiments is that the darker stimuli become more attractive after wing clipping in situations where the animals are faced with a choice of darker and brighter stimuli. One prediction of this hypothesis is that other experiments where the animals face a choice of bright and dark stimuli should also be affected by wing-clipping. To test the generality of the wing-clipping effect and to obtain a third independent test of general activity, we set out to develop a T-maze experiment, where the animals are forced to choose between a dark and a bright arm.

### Wing-clipped flies can show negative photopreference in a T-maze

After several pilot experiments with a variety of different T-maze designs, we arrived at an experimental design where wing-clipped WTB flies would robustly avoid the transparent tube and approach the dark tube (see Material and Methods). As for the BCP, we selected different recovery times (0min, 5min or 24h). Congruent with the BCP results, intact flies showed a positive photopreference, while wing-clipped flies switched to light avoidance and a negative photopreference immediately after their wings were cut (Fig. 3F). These results hold even if the flies are allowed three minutes to choose between the two arms of the T-Maze (Fig. 3G). Also similar to the results in the BCP, we found that the magnitude of the baseline photopreference in intact flies and the wing-clipping effect varied with the genetic background. In the case of the T-Maze, the size of the effect determined whether or not the wing-clipped flies would show positive or negative photopreference (Fig. 3H). Moreover, *CyO* balancer flies also displayed a diminished photopreference, almost an indifference to light, which remained unchanged in wing-clipped animals, in contrast to their siblings (carrying the *tshG80* construct) which showed a clear shift after wing-clipping (Fig. 3I).

### Only manipulations affecting flight-related abilities cause a change in photopreference

While the mutant or transgenic flies used so far may shift their photopreference due to unknown side effects, the shift in wing-clipped flies could in principle be brought about either directly by the injury or indirectly via a detection of flying ability. To distinguish between these two hypotheses, we tested the effects of a series of manipulations (see Materials and Methods, Fig. 1), only some of which affecting some aspect of flight, in BCP and in the T-Maze. First, we evaluated flies with a longitudinal cut through their wings and flies with only one of the two wings completely removed (the side was randomly selected). Both manipulations cause flightlessness. Again, we observed the same shift in photopreference as with standard wing-clipping (Fig. 4A-D). Both flies with longitudinally cut wings (Fig. 4A,B) and one wing removed (Fig. 4C,D) exhibited diminished phototaxis in BCP and a negative photopreference in the T-Maze. During our pilot experiments, we observed that flies with different degrees of injuries on their wings behaved differently. Therefore, we hypothesized that manipulations affecting only some aspects of flight behavior, rather than abolishing flight completely, might lead to less pronounced behavioral changes. Thus, we next compared the behavior of flies whose wings were completely removed, with those where only the tip of the wings had been removed. Flies with partially removed wings are still able to fly, but with reduced torque during turns and reduced lift/thrust [41]. It is worth mentioning that McEwen also attempted to test if the decrease in positive phototaxis was directly proportional to the amount of wing removed, but his low number of replicates, the use of ether as an anesthetic, and his different setup, prompted us to obtain our own data (the same for antenna experiments - see below).

**Figure 4.**
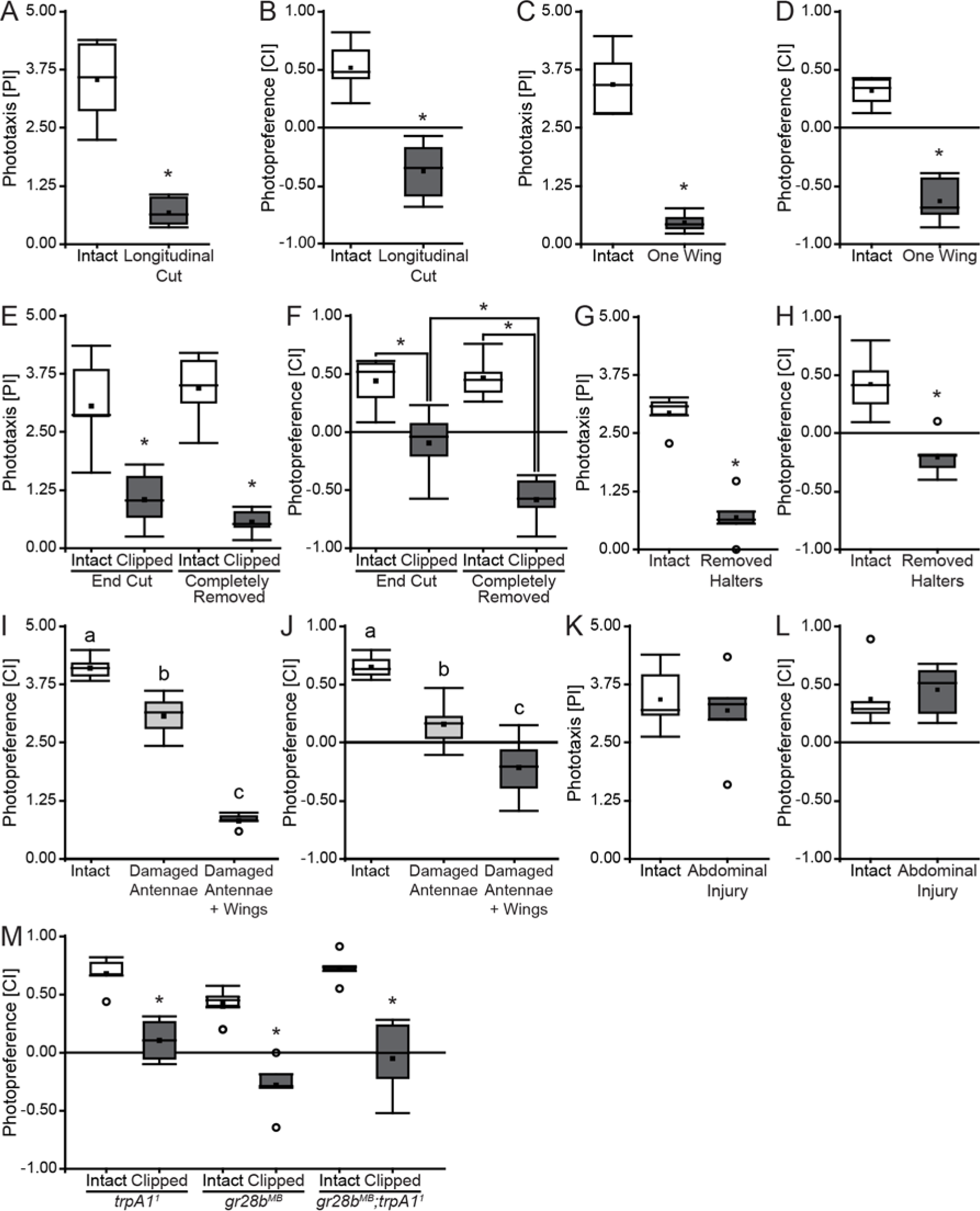
Only flight-affecting manipulations affect photopreference. ***A, C, E, G, I, K***, BCP Performance Index from WTB flies with and without different injuries. ***B, D, F, H, J, L***, T-Maze Choice Index from WTB flies with and without different injuries. ***A, B***, Longitudinal cut of the wings. N=7, *A*: p<0.001, B: p<0.001. ***C, D***, Only one wing cut. N=7, C: p<0.001, D: p<0.001. ***E, F***, Wing clipped at different lengths. Randomized Block Design ANOVA; N=6; *E*: Block p= 0.094, Interaction Wings Integrity (intact or clipped) vs Degree of Injury (completely removed wings or end of the wings cut): p=0.087, Wings Integrity: p<0.001, Degree of Injury: p=0.797; F: Block p= 0.238, Interaction Wings Integrity vs Degree of Injury: p=0.007, simple effects: end cut vs intact: p<0.001, completely removed vs intact: p<0.001, end cut vs completely removed: p<0.001, intact (control from end cut) vs intact (control from completely removed wings): p=0.865. ***G, H***, Both halteres removed. G: N=5, p<0.001, H: N=7, p<0.001. ***I, J***, Both antennae damaged, and both antenna damaged and wings clipped (Damaged Antennae + Wings) I: N=5, ANOVA p<0.001,Tukey’s *post hoc* test (p<0.05; least-significant difference=0.54), J: N=6, ANOVA p<0.001, Tukey’s *post hoc* test (p<0.05; least-significant difference=0.29). Same letter indicates no significant differences. ***K, L***, abdominal wound. K: N=6, p=0.377, L: N=6, p=0.552. ***A, B, C, D, G, H, K, L***, Paired T-Test. M, Thermal sensory deprivation. N=5, T-Test, *trpA1*^*1*^ p<0.001, *gr28b*^*MB*^ p<0.001, *gr28b*^*MB*^;*trpA1*^1^ p=0.001. See figure 1 for detailed graph information.

In both cases (complete and partial removal), injured flies showed a statistically significant reduction in BCP phototaxis and T-Maze photopreference, but both indices were higher in flies with only the end of the wing cut (Fig. 4E,F). In fact, the behavior from both types of injured flies was significantly different from one another in the T-Maze paradigm (Fig. 4F). Therefore, we conclude that behavioral change depends to some extent on the degree of the injury, and on which aspects of flight behavior it affects. To test yet other aspects of flight behavior, we administered injuries that did not affect the wings, in two organs related to flight (halteres and antennae) and one unrelated to flight (the abdomen). In one group of flies, we removed the gyroscopic halteres, mechanosensors involved in sensing body rotation and necessary for free flight [42–45]. In another, we removed the distal segments of the antennae (funiculus and arista), depriving the flies of their most important mechanosensor for airspeed and wind direction [46–48]. The two different treatments both significantly decreased photopreference values (Fig. 4G-J). However, only the manipulation abolishing free flight completely, haltere removal, also led to negative photopreference in the T-Maze (Fig. 4H). Affecting flight stabilization and speed by removing parts of the antennae renders the flies almost indifferent to the light, on average (Fig. 4J). Fully abolishing flight ability in these antenna-damaged flies, yielded negative choice indices (Fig. 4J). Thus, when flies are still able to fly, but individual aspects of flight behavior are disrupted such as stabilization, torque, speed or lift/thrust, their photopreference is less severely affected than when flight is abolished completely. These findings extend the concept of flying ability beyond mere wing utility. To test whether any injury, even one that does not affect any aspect of flight at all, can affect photopreference, we used a small needle to carefully puncture the abdomen of the flies. Consistent with the results so far, a wound in the abdomen did not produce any detectable shift in photopreference (Fig. 4K,L).

### Photopreference shift is not caused by sensory deprivation

A byproduct of manipulations such as cutting the wings or damaging the antennae is the loss of sensory inputs coming from those organs. Therefore, we wondered if any sensory deprivation by itself could cause a dark photopreference in flies which are able to fly. We tested two different thermosensation mutants in the T-Maze paradigm; *trpa1*^*1*^, a long-term thermal preference mutant [49,50], and *gr28b*^*MB*^ which is defective in rapid negative thermotaxis [50]. We also combined *trpa1*^*1*^ and *gr28b*^*MB*^, abolishing thermosensation completely. It is worth to mention that the TrpA1 channel also mediates chemical avoidance via gustatory neurons [51,52], and Gr28b is expressed in HC-neurons located in the same portion of the antennae damaged with our manipulation [50]. The wings-intact mutants all showed a positive photopreference (Fig. 4M), indicating that photopreference is not automatically affected when any sensory modality is knocked out. Corroborating this observation was a sharp drop in photopreference when the wings were clipped in these mutants (Fig. 4M).

### The shift in photopreference is reversible and traces wing utility

If flies were monitoring the different aspects of their flying abilities and changing their photopreference accordingly, one would expect that transient impairments in wing utility would cause transient changes in photopreference. To examine the reversibility of the behavioral shift, we designed two complementary experiments. In the first, we tested *WTB* flies in BCP and T-Maze before and after gluing, as well as after ungluing their wings. Wing gluing perfectly reproduced the wing-clipping effect, evidenced by a clear reduction of the PI and CI (Fig. 5A,B), showing again that the shift in photopreference is independent from the cause of the flightlessness. Remarkably, normal photopreference was restored after cleaning the wings of the tested flies (Fig. 5A,B).

**Figure 5.**
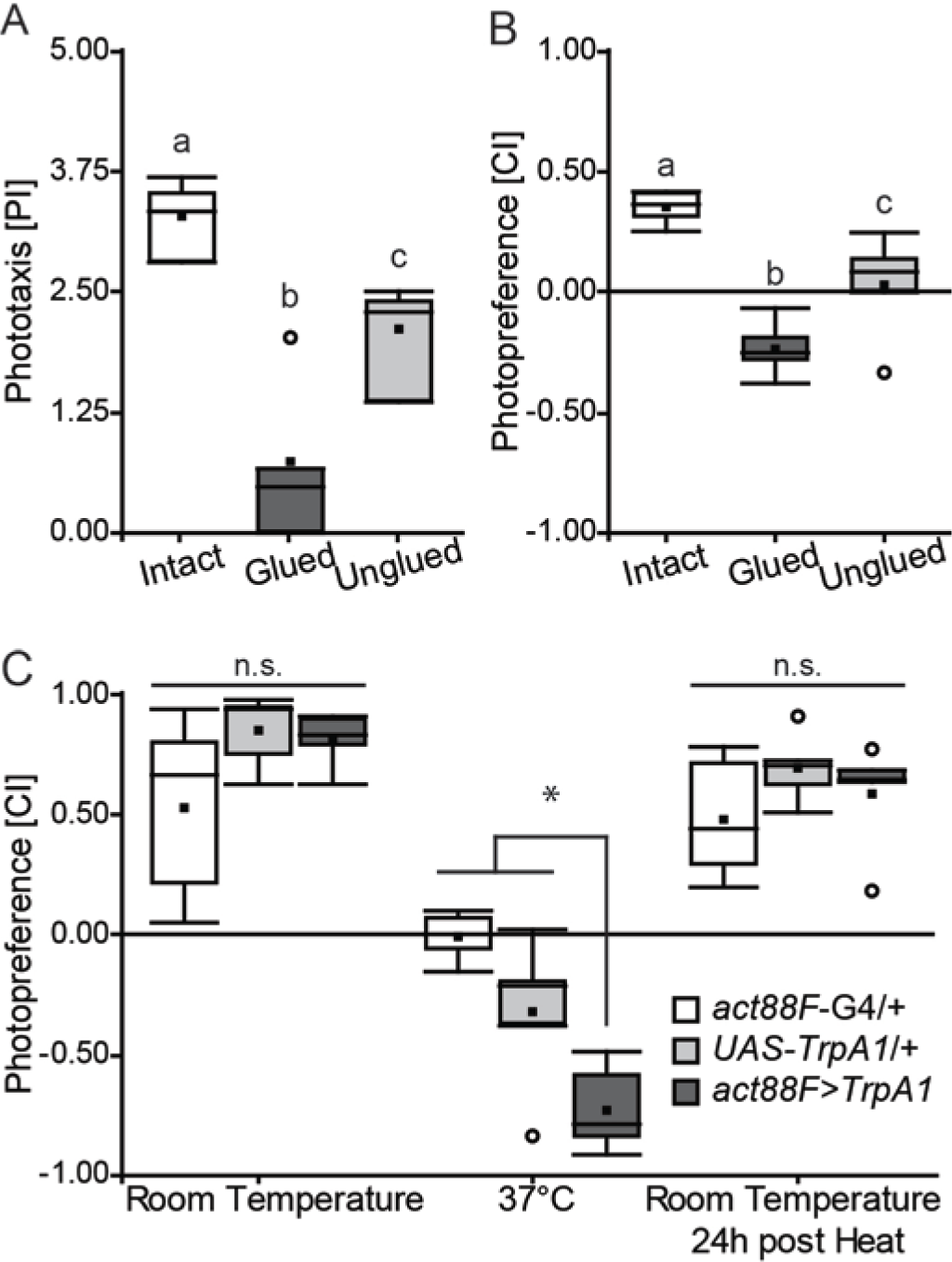
Photopreference changes together with wing utility in a reversible manner. **A**, BCP tests in flies before, during and after their wings had been rendered useless by applying (and then removing) sucrose solution. Randomized Block Design ANOVA, N=4, Block p=0.091, ANOVA p<0.001, Tukey’s post hoc test (p<0.05; least-significant difference=1.0257). ***B***, T-Maze. Randomized Block Design ANOVA, N=5, Block p=0.173, ANOVA p<0.001, Tukey’s post hoc test (p<0.05; least-significant difference=0.232). Same letter indicates no significant differences. ***C***, Genetic manipulation of IFM contraction and wing utility. T-Maze Choice Index before, during and after 37°C exposure of experimental and control flies. Randomized Block Design ANOVA, N=5, Block p=0.152, Interaction Genotype vs Temperature: p<0.001, simple effects with Tukey’s *post hoc* test (p<0.05): least-significant difference=0.349, Room Temperature: p=0.073, 37°C: p<0.001, Room Temperature 24h post heat: p=0.344. * indicates significant differences, n.s. means not significant. See figure 1 for detailed graph information.

In our complementary approach, we manipulated wing utility by reversibly altering Indirect Flight Muscle (IFM) contraction, expressing the temperature-sensitive *TrpA1* channel under the promoter of the IFM-specific gene *actin 88F (act88F)*, using the *act88F-GAL4* [53] driver. At room temperature, experimental flies tested in our T-Maze were indistinguishable from their genetic controls. However, at 37°C, when *TrpA1* caused a sustained IFM contraction disrupting wing movements, the same flies showed a marked preference for the dark arm of the maze that fully recovered when they were tested back at room temperature on the following day (Fig. 5C). The genetic controls also showed a CI decrease at 37°C, but it was less pronounced and significantly different from the experimental group. In sum, these results show that flies adjust their photopreference in accordance with their wing utility. Moreover, these changes are immediate and reversible.

### Wing-clipping effect is not dependent on known learning and memory processes

The reversibility of the shift in photopreference is reminiscent of a learning process where the animal may evaluate its flight capabilities at one point and then remember this outcome until the next evaluation. For instance, the animals may attempt flight and immediately learn about the futility of their attempt. Until the next attempt, the flies remember this state and shift their photopreference accordingly. To test this hypothesis, we screened a selection of mutant/transgenic fly lines with a variety of known learning and memory impairments using BCP. We selected lines known to affect classical olfactory conditioning/operant world-learning, operant self-learning, or any Mushroom Body-dependent learning processes. In order to avoid differences related to specific locomotor characteristics from the different lines, here again the wing-clipping effect was assessed with the Effect Size. None of the lines tested showed any wing-clipping effect at all. All lines showed a clear behavioral change after wing-clipping, evidenced by a decrease in their PI with and *Effect Size* around -0.6 or more, irrespective of the baseline value (Fig. 6A, B).

**Figure 6.**
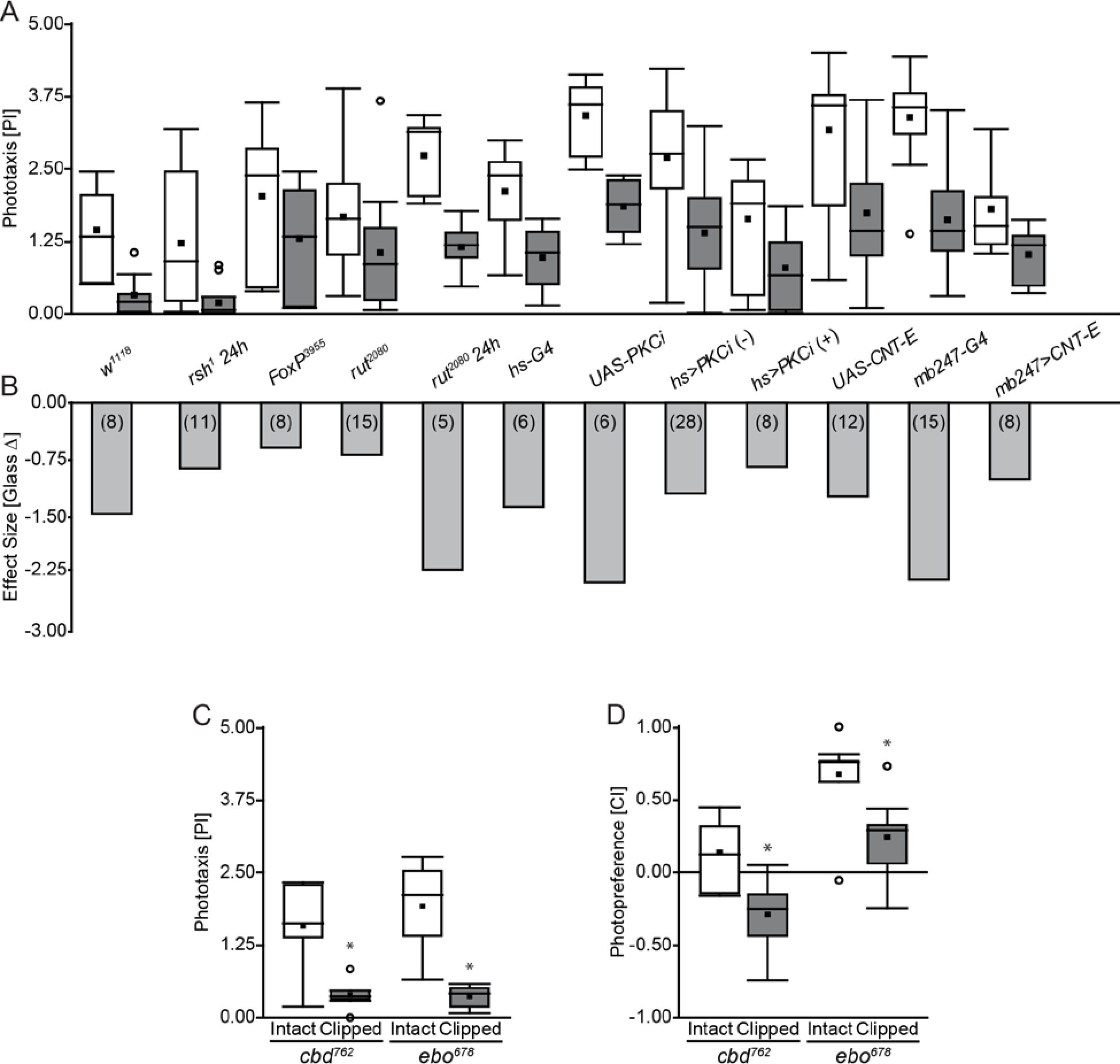
The wing-clipping effect is independent from known learning/480 memory processes or neuropil areas associated with learning. **A**, Performance Index of mutants and transgenic flies with learning and memory impairments, before and after clipping their wings. **B**, Effect Size of wing clipping on BCP for several lines with learning and memory impairments and their controls. N=Numbers in brackets. **C**, **D**, Behavioral performance from two structural Central Complex mutants with intact and clipped wings on BCP (c) and T-Maze (d). Paired T-Test. c, cbd^762^, N= 6, p=0.005; ebo^678^, N= 6, p=0.004. d, cbd^762^, N= 8, p=0.002, ebo^678^, N= 7, p<0.001. See figure 1 for detailed graph information.

### The behavioral switch is not central complex-dependent

The central complex is a higher-order neuropil related to locomotion [54,55], visual information processing [56], orientation [57], visual pattern recognition [58,59] and spatial working memory [60]. As many of these functions may be important for either phototaxis or its flexibility, we tested two structural mutants of this neuropil, Central Body Defect (*cbd*^762^) and Ellipsoid Body Open (*ebo*^678^). However, wing-clipped *cbd*^*762*^ as well as *ebo*^*678*^ flies both showed a clear significant change in their photopreference measured either in BCP or T-Maze (Fig. 6C,D). We note that, although *ebo*^*678*^ wingless flies still showed a preference for the bright tube in the T-Maze, their PI was significantly decreased in comparison with intact *ebo*^*678*^ flies. While more sophisticated manipulations of central complex function are clearly warranted, we tentatively conclude that if the central complex plays a role in this process, it is likely not a crucial one, or one that does not require an anatomically intact central complex.

### DA and OA differently modulate intact and wingless fly behavior

In the absence of any evidence that any of the known learning processes or neuropils known to be relevant for learning or other aspects of orientation/choice behaviors are crucial for the shift in photopreference, we explored the hypothesis that any unknown learning mechanism as well as an unknown constant monitoring of flying ability may rely on a re-valuation of sensory input after wing manipulation. That is, whether or not any memory is involved, the consequence of being rendered flightless may be identical: a re-valuation of sensory input, such that previously attractive stimuli become more aversive and previously aversive stimuli become more attractive. Biogenic amines have long been known for their role in mediating the processing and assignment of value [4,9,11–13,15,21,61–67]. If indeed it is the photopreference that is shifted when a fly’s flying ability is altered, it is straightforward to hypothesize that the two biogenic amines most known for being involved in valuation in *Drosophila*, octopamine (OA) and dopamine (DA), may be involved in this instance of value-based decision-making as well. Moreover, mutant flies that lack tyrosine hydroxylase (th) only in the nervous system, i.e. neuronal specific DA-deficients, show reduced phototaxis in BCP [66] further motivating the manipulation of this amine pathway. Finally, flies without OA show a pronounced impairment in flight performance and maintenance [68], making OA an interesting candidate for testing photopreference as well.

To evaluate the involvement of DA and OA neurons for photopreference, we acutely disrupted synaptic output from two separate groups of neurons by expressing the temperature-sensitive form of dynamin (*Shibire; shi*^*TS*^, [69]) either under control of the th-GAL4 driver (driving in dopaminergic neurons) or under control of the *tdc2-GAL4* driver (driving in octopaminergic, as well as tyraminergic, neurons). We tested the resulting transgenic flies with and without wings in BCP and T-Maze. Although BCP and T-Maze results tended to agree, we only obtained clear results in our T-Maze experiments. The reason for the less clear results in the BCP was a genotype-independent and long-lasting effect of the temperature switch on the flies’ PI in the BCP. Hence, we show the results from the T-Maze experiments here and the BCP results are available for download with the rest of the raw data. In the T-Maze at permissive room temperature, when dynamin is in its wild type conformation, in all tested groups, flies with intact wings showed positive CIs, while wing-clipped flies showed negative CIs (Fig. 7A,B). In contrast, when the same experiment was performed at the restrictive 32°C (i.e., blocking synaptic activity), we found opposite effects in flies with dopaminergic, and octopaminergic/tyraminergic neurons blocked, respectively. While disrupting synaptic output from dopaminergic neurons appeared to have little if any effect on clipped animals, flies with intact wings shifted their preference to the dark tube (Fig. 7A), rendering their CI indistinguishable from that of their wingless siblings with which they were tested (Fig. 7B). Conversely, blocking synaptic output from octopaminergic neurons only affected wingless flies, which now preferred the bright arm of the maze (Fig. 7B), similar to their siblings capable of flight with which they were tested (Fig. 7A). Replicating the reversibility described above, after a 24h recovery phase, flies tested at room temperature showed wild type behavior, meaning positive photopreference for intact flies and negative photopreference for wing-clipped flies (Fig. 7A,B). The conventional interpretation of these results is that synaptic transmission from octopaminergic/tyraminergic (OA/TA) neurons is necessary for shifting the photopreference towards darkness in flightless flies, while synaptic transmission from DA neurons is necessary for setting the preference of intact flies towards the bright arm.

**Figure 7.**
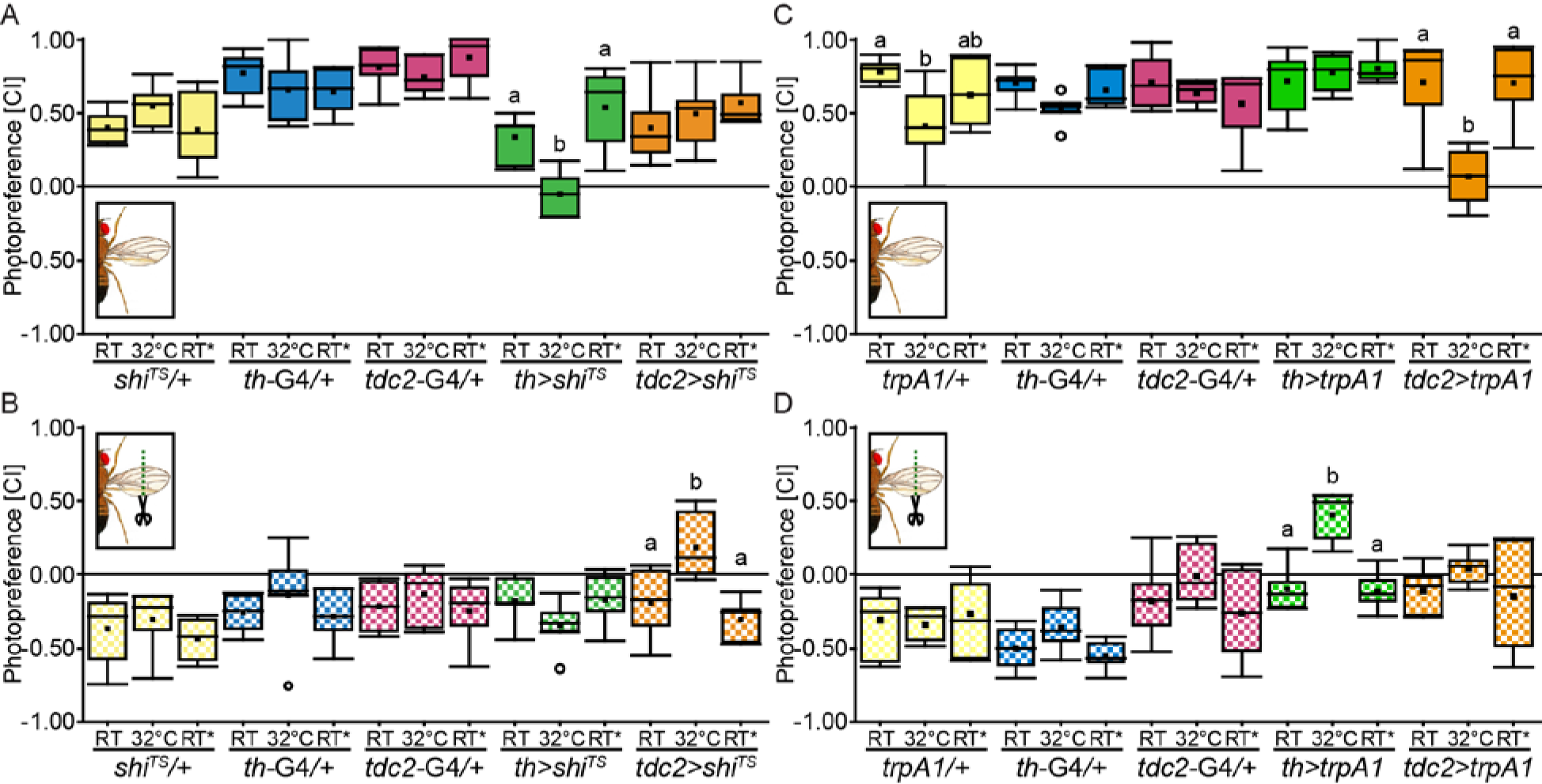
Dopamine and Octopamine are necessary and sufficient to modulate phototactic behavior, but with opposite effects. ***A, B***, Photopreference from flies with (A) and without (B) wings before, during and after DA or OA/TA neuron silencing. **A,** Randomized Block Design ANOVA, Block p=0.026, Interaction Genotype vs Temperature p<0.001, simple effects with Tukey’s *post hoc* test (p<0.05, least-significant difference=0.24, *tdc2>shi*^*-ts*^ least-significant difference= 0.263): *shi*^*-ts*^/+ p=0.208, th-GAL4/+ p=0.417, tdc2-GAL4/+ p=0.428, *th>shf*^*s*^ p<0.001, *tdc2>sh(*^*s*^ p=0.242. N=6 except for *tdc2>sh(*^*s*^ RT* (N=5). **B**, Randomized Block Design ANOVA, Block p=0.006, Interaction Genotype vs Temperature p=0.02, simple effects with Tukey’s *post hoc* test (p<0.05, least-significant difference=0.278, *tdc2>shi* least-significant difference= 0.288): *shi*^*-ts*^/+ p=0.533, th-GAL4/+ p=0.394, tdc2-GAL4/+ p=0.6, *th>shi*^*ts*^ p=0.262, *tdc2>shf*^*s*^ p<0.001. N=6 except for *tdc2>shi*^*-ts*^ RT* (N=5). ***C, D***, Photopreference from flies with (C) and without (D) wings before, during and after DA or OA neuron activation. ***C***, Kruskal-Wallis for temperature factor comparison within genotypes (alpha after correction=0.013): *trpA1/+* p=0.012, *th-*GAL4/+ p=0.069, tdc2-GAL4/+ p=0.667, *th>trpA1* p=0.97, *tdc2>trpA1* p=0.004. th-GAL4/+ and *th>trpA1,* N=6; *trpA1/+*, tdc2-GAL4/+ *tdc2>trpA1*, N=7. *D*, Kruskal-Wallis for temperature factor comparison within genotypes (alpha after correction=0.013): *trpA1/+* p=0.834, th-GAL4/+ p=0.15, tdc2-GAL4/+ p=0.126, *th>trpA1* p=0.005, *tdc2>trpA1* p=0.415. th-GAL4/+ and *th>trpA1*, N=6; *trpA1/+*, tdc2-GAL4/+ *tdc2>trpA1*, N=7. Different letters indicate significant differences between temperatures for each genotype (only shown for genotypes where the factor temperature had a statistically significant effect). See figure 2 for detailed graph information.

We also transiently activated OA/TA and DA neurons, respectively, using the temperature sensitive *TrpA1* channel [49], while testing the flies for their photopreference. Again, at room temperature, when the channel is closed, flies with and without wings behaved similar to wild type animals (Fig. 7C,D). However, when tested in the same experiment at 32°C, where the *TrpA1* channel is open and depolarizes the neurons in which it is expressed, the flies showed a change in their behavior. Flies with clipped wings and activated DA neurons now preferred the bright arm of the maze, with no effect on intact flies (Fig. 7D). Conversely, activating OA/TA neurons only had an effect on flies with intact wings, abolishing their previous preference for the bright arm of the maze (Fig. 7C), rendering them indistinguishable from their wingless siblings with which they were tested, but which did not show any significant effect (Fig. 7D). Again, when tested back at room temperature 24h later, wild type behavior was restored. The conventional interpretation of these results is that active OA/TA neurons are sufficient for shifting photopreference towards the dark arm of the maze, while the activation of DA neurons is sufficient to set the flies’ preference towards brightness.

## Discussion

McEwen’s discovery captured our attention because of its implications for the supposed rigidity of simple behaviors. We first reproduced the findings of McEwen [33] and Benzer [32] that wing manipulation leads to a decrease in *Drosophila* phototaxis (Fig. 2). Slightly altering the conditions of the BCP and comparing performance between two additional experiments, we found that the decrease in phototaxis is not due to hypoactivity of wing-manipulated flies, but to a more general change in the flies’ assessment of their environment (Fig. 3). We discovered evidence that the BCP is just one of several experiments that can measure a fly’s general photopreference. Manipulating the wings modulated this preference in all of the selected experiments such that compromised wing utility yielded a decreased preference for brightness (bright stimuli) and an increased preference for darkness (dark stimuli) across the experiments chosen (Fig. 3). However, of these experiments, only the BCP can be argued to test phototaxis proper. In Buridan’s Paradigm the flies walk between two unreachable black stripes; and in the T-Maze, the flies choose between a dark tube and a bright one where the light is coming from an angle perpendicular to their trajectory. Neither of the two paradigms is testing taxis to or away from a light source. Interestingly, in our pilot experiments, we have tested phototaxis in different variations of the T-maze with various LEDs placed at the end of one of two opaque tubes and only found a reduction of phototaxis and never negative phototaxis (unpublished observation). In fact, in these pilot experiments we have observed every possible difference between flying and manipulated flies. In the end, we chose the experimental design that yielded positive and negative scores, respectively, in WTB flies purely for practical reasons. Other wild type strains, such as some Canton S substrains, do not show a negative photopreference in the T-Maze after wing clipping (Fig. 3H). Taken together, these lines of evidence strongly suggest that photopreference in *Drosophila* is a strain-specific continuum where experimental design assigns more or less arbitrary values along the spectrum. In some special cases, this photopreference manifests itself as phototaxis. If that were the case, phototaxis would constitute an example of a class of experiments not entailing a class of behaviors.

This insight entails that manipulations of different aspects of flight ought to affect this continuum in different ways. Complete loss of flight ought to have more severe effects than manipulations affecting merely individual aspects of flight behavior, such as wing beat amplitude/frequency (i.e., lift/thrust), torque, flight initiation, flight maintenance, proprioception or motion/wind-speed sensation. We have found some evidence to support this expectation. For instance, clipping only the tips of the wings does not eliminate flight, but affects torque as well as lift/thrust [41,70]. Flies with the tips of their wings cut behaved indifferently in the T-Maze and do not avoid the bright tube (Fig. 4F). Flies without antennae are reluctant to fly and have lost their main sense of air speed detection [46–48], but they are still able to fly. Also these flies do not become light averse in the T-Maze after the manipulation, but indifferent. Only clipping the wings in these flies abolishes their flight capabilities completely and yields negative scores (Fig. 4I). Flies with removed gyroscopic halteres, on the other hand, are severely affected in their detection of rotations and usually do not fly [42–44], despite being able to still beat their wings and control flight direction using vision alone in stationary flight [42,43]. These flies avoid the bright arm of the T-Maze. Finally, injuries to flight-unrelated parts of the fly’s body did not affect photopreference (Fig. 4K, L), ruling out the preference of darkness being a direct escape response due to bodily harm. Further research is required to establish a quantitative link between the many different aspects of flight behavior and their relation to photopreference.

Taken together, our experiments so far demonstrate that 1) the physical state of the wings with regard to their shape, form or degree of intactness influences photopreference (Figs. 2-4). 2) The capability to not just move the wings, but specifically to move them in a way that would support flight (Figs. 2, 3, 5) also influences the flies’ photopreference. 3) The state of sensory organs related to flight such as antennae or halteres also exerts such an influence, while non-flight-related sensory deprivation shows no such consequences (Fig. 4). This multitude of flight-related aspects extends the concept of flying ability beyond mere wing utility: manipulating seemingly any aspect of the entire sensorimotor complex of flight will affect photopreference, and do so reversibly (Fig. 5). As it appears that any aspect of flight, sensory or motor, is acutely linked to photopreference, it is straightforward to subsume all of these aspects under the term ‘flying ability’, emphasizing that flying ability encompasses several more factors in addition to wing utility. The observation that each fly, when it is freshly eclosed from the pupal case and the wings are not yet expanded, goes through a phase of reduced phototaxis that extends beyond wing expansion until the stage when its wings render it capable of flying [71], lends immediate ethological value to a neuronal mechanism linking flying ability with photopreference.

One possibility how the link between flying ability and photopreference may be established mechanistically is via a process reminiscent of learning: at one time point, the flies register a sensory or motor deficit in their flight system and at a later time point, they use this experience when making a decision that does not involve flying. Once flying ability is restored, the same choice situation is solved with a different decision again in the absence of flight behavior. How the flies accomplish this learning task, if indeed learning is involved, is yet unknown, but we tentatively conclude that it is unlikely that any of the known learning pathways or areas involved in different forms of learning play more than a contributing role (Fig. 6). While the molecular learning mechanism remains unidentified, the process appears to be (near) instantaneous (Figs. 2, 3). Even though we cannot rule out that an unknown learning mechanism exists which is unaccounted for in our screen, we conclude that at least none of the known learning mechanisms suffices to explain the complete effect size of the shift in photopreference. These results corroborate the findings above, that the switch is instantaneous and does not require thorough training or learning from repeated attempts to fly, let alone flight bouts. They do not rule out smaller contributions due to these known learning processes or an unknown, fast, episodic-like learning process. It is also possible, that the flies constantly monitor their flying ability and hence do not have to remember their flight status. Despite these ambiguities, we have been able to elucidate some of the underlying neurobiological mechanisms. Much as in other forms of insect learning and valuation [72–76], neurons expressing the biogenic amine neuromodulators OA and DA appear to have opposite functions in the modulation of photopreference (Fig. 7).

Although both DA and OA play some role in different aspects of flight behavior [68,77–79], these cannot explain our results. In general, our biogenic amine neuron manipulated flies escape their vial via flight if granted the opportunity. Thus, flight is not abolished in any of our transgenic lines affecting OA, TA or DA neurons. However, there may be more subtle deficits in less readily perceived aspects of flight. Experiments performed with mutant flies lacking OA demonstrated that OA is necessary for initiation and maintenance of flight [68]. However, in our paradigm, silencing OA/TA neurons promoted approaching light, the opposite effect of what would be expected for a flightless fly (Fig. 7 B). Activating these OA/TA neurons, however, rendered the flies indifferent in the T-Maze. OA/TA appear to be involved in flight initiation and maintenance via opponent processes [68]. Transient activation of OA/TA neurons may lead to a subtle alteration of flight performance and reduce photopreference in these flies. Similarly, it has been shown that altering the development of specific DA neurons results in flight deficits (reduction of flight time or loss of flight, depending on the treatment [78,79]). Our manipulations lasted for approximately 30 min during adulthood, ruling out such developmental defects. Work in the laboratory of Gaiti Hasan has also found that silencing of three identified TH-positive interneurons for several days in the adult animal compromises flight to some extent (wing coordination defects during flight initiation and cessation [77]). Our much shorter manipulation does not lead to any readily observable flight defect. However, one needs not discuss whether or not our aminergic manipulations may have had subtle effects on some aspects of flight behavior, as we can compare these flies to the wing-clipped siblings with which they were tested simultaneously (i.e., the flies with the maximum shift in photopreference due to completely abolished flight). Comparing the intact DA-inactivated flies and OA/TA-activated flies (Fig. 7 A,C) with their respective wingless siblings (Fig. 7 B,D) reveals that the CIs of the pairs of groups become essentially indistinguishable at the restrictive temperature. In other words, intact flies where DA neurons have been inactivated or OA/TA neurons have been activated behave as if their wings had been clipped and their flight capabilities abolished completely, despite them being capable of at least some aspects of flight. Hence, even if there were some contribution of some aspect of flight behavior being subtly affected by manipulating these aminergic neurons, there is a contribution of activity in these neurons that goes beyond these hypothetical flight deficits. Therefore, we conclude that neither the OA/TA, nor the DA effects can be explained only by subtle defects in one or the other aspect of flight behavior in the manipulated flies.

The precise neurobiological consequences of manipulating OA/TA and DA neurons, respectively, are less certain, however. Not only are the two driver lines *(th-GAL4* and *tdc2-*GAL4) only imperfectly mimicking the expression patterns of the genes from which they were derived. Our effectors, moreover, only manipulated the activity of the labeled neurons. One manipulation (*shi*^*TS*^) prevents vesicle recycling and likely affects different vesicle pools differentially, depending on their respective release probabilities and recycling rates. The other effector *(TrpA1)* depolarizes neurons. It is commonly not known if the labelled neurons may not be co-releasing several different transmitters and/or modulators in the case of supra-threshold depolarization. Hence, without further research, we can only state the involvement of the labelled neurons, which as populations are likely to be distinct mainly by containing either DA or OA/TA, respectively. If it is indeed the release of these biogenic amines or rather the (co-)release of yet unknown factors in these neuronal populations remains to be discovered. Further research will also elucidate the exact relationship between the activities of these two neuronal populations and whether/how it shifts after manipulations of flying ability (Fig. 8).

**Figure 8.**
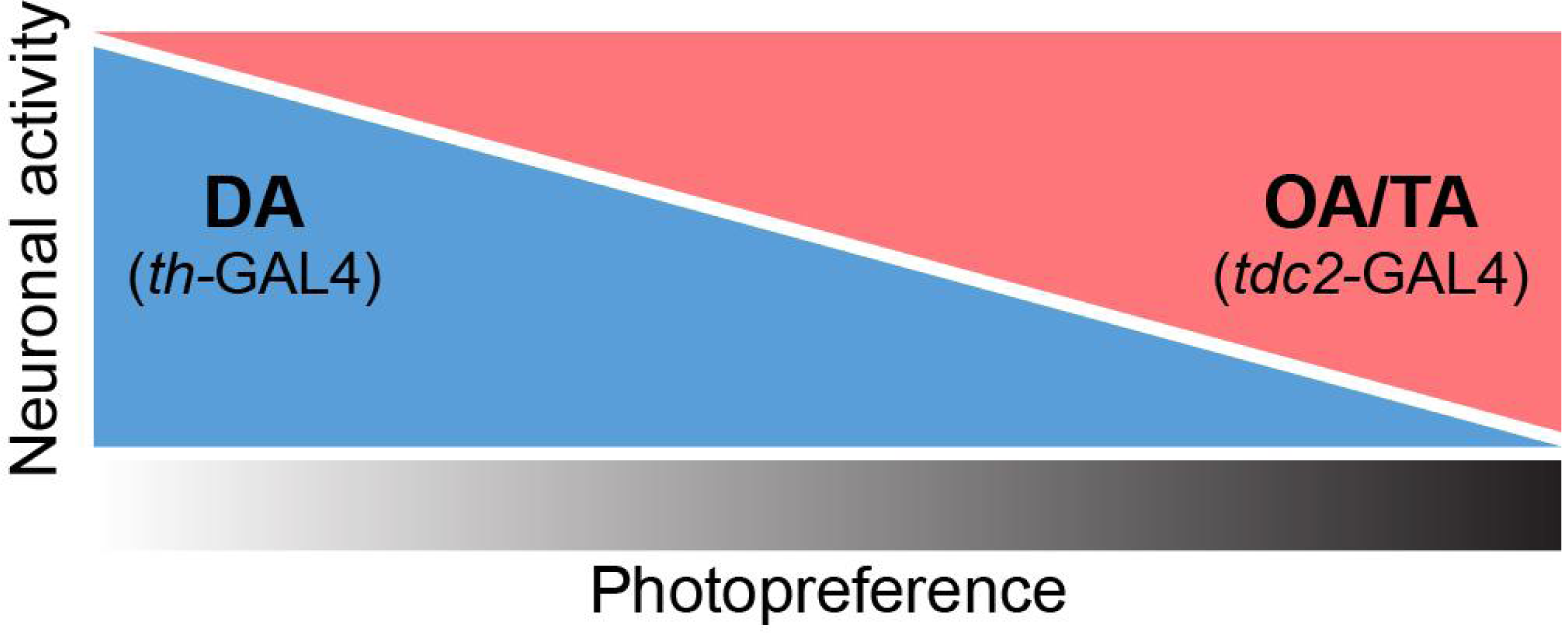
Schematic illustration of the potential dependence of photopreference on the activity of aminergic neurons. Depending on several factors (e.g., the status of its flight apparatus), individual flies may fall anywhere on the photopreference spectrum (grayscale): approaching light, avoiding it or behaving indifferently. Increasing neuronal activity in tdc2-GAL4 positive neurons (red) or decreasing neuronal activity in *th-*GAL4 positive neurons (blue), each alone promoted a preference of darkness (shift to the right of the spectrum) in flies able to fly, which normally prefer brightness over darkness. In contrast, increasing neuronal activity in th-GAL4 neurons (blue) or decreasing neuronal activity in tdc2-GAL4 neurons (red), each alone promoted preference of brightness (shift to the left of the spectrum) in wing-clipped flies, which normally tend to avoid brightness. It is straightforward to hypothesize that the quantitative relationship between two opponent processes (potentially based on OA/TA and DA action) constitutes one mechanism mediating photopreference in *Drosophila*. In this figure, we depicted this relationship as linear for illustrational purposes only.

In conclusion, our findings provide further evidence that even innate preferences, such as those expressed in classic phototaxis experiments, are not completely hard-wired, but depend on the animal’s state and presumably other factors, much like in the more complex behaviors previously studied [21–26]. This endows the animal with the possibility to decide, for example, when it is better to move towards the light or hide in the shadows. Moreover, the fact that flies adapt their photopreference in accordance with their flying ability raises the tantalizing possibility that flies may have the cognitive tools required to evaluate the capability to perform an action and to let that evaluation impact other actions - an observation reminiscent of meta-cognition.

## Acknowledgments

We would like to thank the Bloomington Stock Center and the colleagues listed in the Methods section for sharing fly stocks. We would like to thank former undergraduate students Lidia Castro, Ben Beuster, Marc-Nicolas Rentinck, Christian Rohrsen, Lucie Dietrich, Christin Dröger, Chang Hae In, Asli Akin and Malte Hahn for technical assistance and (pilot) data collection. We would also like to thank N. Muraro and L. Frenkel for critical reading of the manuscript. EAG was supported by a DAAD research scholarship.

## Author’s contribution

EAG, JC and BB designed the experiments, EAG and JC performed the experiments, EAG and BB wrote the manuscript.

